# Tachykinin 1-expressing neurons in the lateral habenula signal negative reward prediction error

**DOI:** 10.1101/2025.10.11.680558

**Authors:** Kana E Suzuki, Tharusha A Seagoe, Blake Holcomb, Jackie R Kuyat, Emily L Sylwestrak

**Affiliations:** Department of Biology, University of Oregon, Eugene, OR; Institute of Neuroscience, University of Oregon, Eugene, OR

## Abstract

Evaluating outcomes to accurately predict which actions lead to reward is crucial for survival. Discrepancies between expected and realized outcomes, termed reward prediction errors (RPEs), serve as a teaching signal to update subsequent predictions and promote adaptive behavior. Neural correlates of RPEs have been identified in several brain regions, including the lateral habenula (LHb), which contains a subpopulation of neurons encoding negative reward prediction error (nRPE) that are excited by worse-than-expected outcomes and inhibited by better-than-expected outcomes. LHb projections to the midbrain shape firing in dopaminergic neurons and play a well-established role in reward learning and decision-making. However, the LHb engages in a wide variety of behaviors beyond reward processing, and it remains unclear whether these diverse functions are mediated by specific transcriptionally defined cell types. Little is known about the transcriptomic identity of nRPE-encoding neurons, limiting our understanding of the specific role of these signals in outcome valuation. Using cell type-specific recording in mice performing reward-guided tasks, we demonstrate neurons expressing the neuropeptide gene *Tachykinin-1* (*Tac1*) represent a subpopulation of LHb neurons that encode nRPE. We found LHb^Tac1^ activity is sensitive to changes in both the expected value and realized value of rewards, and scales with the magnitude of the difference. Further, LHb^Tac1^ neurons show little modulation to other task-related events, or to innately aversive stimuli that engage a broader population of LHb cell types. Together, these data demonstrate that *Tac1* marks a subpopulation of LHb neurons that preferentially encodes nRPE. Our results provide insight into cell type-specific contributions of habenular neurons in nRPE signaling and open avenues for more targeted manipulations of nRPE-encoding neurons to understand their role in reward-guided behavior.

## INTRODUCTION

To maximize reward in dynamic environments, animals must learn which actions lead to positive outcomes and rapidly adapt when those actions no longer yield rewards. This process relies on predicting likely outcomes, evaluating the results, and adjusting behavioral strategies accordingly. Classical models of reward learning posit that updates occur at mismatches between predictions and outcomes, called reward prediction errors^1–4^. Rewards that are unexpected—or unexpectedly good—produce a positive prediction error (pRPE), whereas reward outcomes that fail to meet expectations produce a negative prediction error (nRPE). Dopaminergic (DA) neurons in the ventral tegmental area (VTA) of monkeys^4^ and rodents^5– 8^ signal RPEs, displaying increased firing at better-than-expected outcomes and decreased firing at worse-than-expected outcomes. Similar correlates of RPE have been shown in the nucleus accumbens^9,10^ and ventral pallidum^11^ in multiple species^7,12–14^. A subpopulation of neurons in LHb also encodes RPEs but in an inverted manner to DA neurons: they are *excited* by negative RPE and *inhibited* by positive RPE^15–20^. The LHb provides disynaptic inhibitory input to dopaminergic neurons in the VTA via GABAergic neurons in the rostromedial tegmental nucleus (RMTg) which invert the signal^15,21–25^. LHb lesions degrade RPE signals in the VTA, supporting a model where LHb→RMTg→VTA signaling shapes dopaminergic RPE^26,27^. However, the LHb has also been implicated in a wide variety of other behaviors including aversive learning^28–30^, anxiety^31^, stress^20,32^, attention^33^, innate defensive strategies^30,34^, social behaviors^35,36^, aggression^37–39^, and parental behavior^40,41^. RPE-encoding neurons account for only a subset of LHb neurons, but their molecular identity is unknown. Their diverse functional roles pose a significant barrier to interpreting causal manipulations that affect all LHb neurons. Transcriptional analyses indicate a variety of molecularly distinct cell types that could serve as a substrate to segregate information from diverse behavioral streams^42–46^, but data linking cell types to behaviors remains sparse^39,47^. The gene *Tac1* is an established cell type marker in the medial habenula, where it marks reward-responsive neurons^42–46,48^. It is also expressed by a subpopulation of neurons distributed throughout the LHb (Figure 1A, S1A) that have been shown to respond to reward outcomes^48^, prompting us to hypothesize that they may encode RPEs. Using cell-type specific fiber photometry recordings in freely moving mice across multiple reward learning paradigms and aversive behavioral contexts, we show that LHb^Tac1^ neurons track changes to task variables that alter both the predicted and realized value of reward and their activity scales with the magnitude of the prediction-outcome mismatch—hallmarks of a neural representation of reward prediction error.

**Figure 1.**
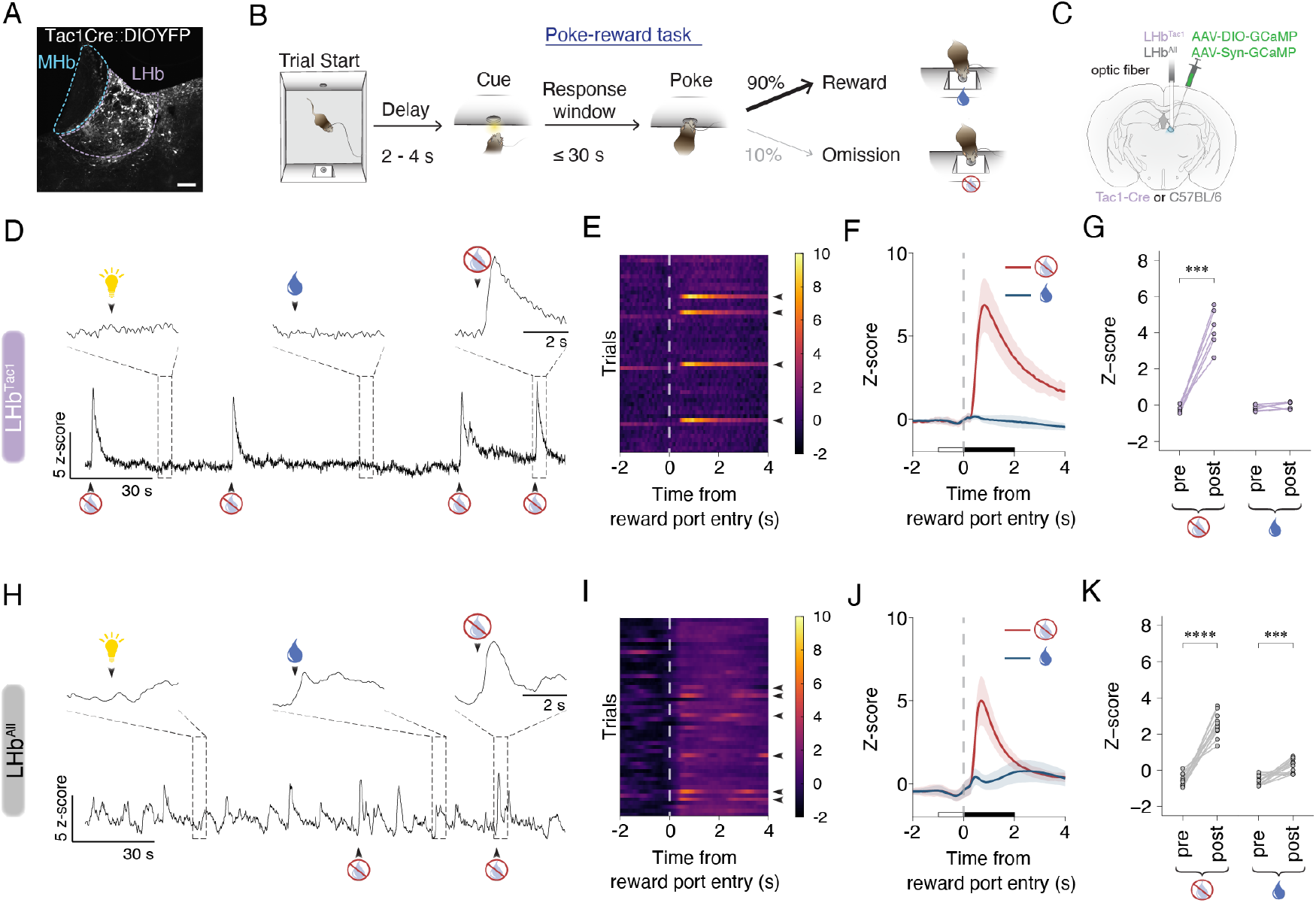
LHb^Tac1^ neurons preferentially respond to reward omissions. (A) Example image of AAV-Ef1𝔞-DIO-YFP virus injection into lateral habenula of *Tac1*-Cre mice. Scale bar = 100 µm (B) Schematic of poke-reward task. Nose poke in illuminated port triggers sucrose water delivery in port on the opposite end of chamber. In 10% of trials, no sucrose water reward is delivered. Premature pokes in cue port result in 10 s timeout. (C) Schematic of AAV-DIO-GCaMP or AAV-Syn-GCaMP injection and optic fiber placement in lateral habenula of *Tac1*-Cre or C57BL/6 mice. (D – K) Neural responses to behavioral stimuli were recorded in LHb^Tac1^ (D-G) or LHb^All^ (H-K) populations using fiber photometry. (D, H) Example trace from fiber photometry recordings during poke-reward task, z-scored across the session. Callouts indicate 5 s activity window around onset of cue (left), reward consumption (middle), and reward omission (right). (E, I) Example of 50 correct trials in a fiber photometry session with 10% of rewards omitted. Data are aligned to head entry into reward port. Pseudocolor denotes z-score. Arrowheads indicate trials with omitted rewards. (F, J) Averaged responses aligned to reward omission (red) and reward consumption (blue). (G, K) Average fluorescence before and after reward omission (left) and reward consumption (right) for each animal, calculated in a 1 second window before the reward onset (open bar) and 2 seconds after the reward onset (filled bar), respectively. Paired t-test with FDR correction for multiple comparisons. LHb^Tac1^: omission, p < 0.001; reward, p = 0.24. LHb^All^: omission, p < 0.0001; reward, p < 0.001. *** p< 0.001, **** p < 0.0001. LHb^Tac1^, n = 6; LHb^All^, n = 12. Error bars indicate standard deviation.

## RESULTS

### LHb^Tac1^ neurons are preferentially engaged during reward omissions

To investigate how LHb^Tac1^ neurons encode reward-related information, we used fiber photometry in freely moving mice performing a simple “poke-reward” operant task. Water-restricted mice were trained to nose poke into an illuminated initiation port on one side of an operant chamber. Successful pokes resulted in delivery of a sucrose water reward on the other side of the chamber (Figure 1B). Trained mice showed shorter latencies to nose poke in the initiation port, and to collect reward following water delivery (Figure S1B-D) indicating they had learned the task. To monitor reward-related activity in LHb^Tac1^ neurons, we injected AAV1-DIO-GCaMP6 into the LHb of *Tac1*-Cre mice and implanted an optical fiber over the injection site (Figure 1C). We recorded the activity of LHb^Tac1^ neurons while mice performed the poke-reward task with a probabilistic reward schedule: rewards were delivered in 90% of trials and omitted in 10% of trials. LHb^Tac1^ neurons showed responses to reward omissions but not reward delivery (Figure 1D-G). Unlike the dynamic responses at reward omission, LHb^Tac1^ neurons displayed very little modulation outside of that period (Figure 1D), with no significant changes during other behavioral events in the task (e.g. cue, nosepoke) (Figure S1E-G). To compare LHb^Tac1^ activity to pan-neuronal LHb activity, we injected AAV1-Syn-GCaMP in C57BL/6 mice performing the poke-reward task (LHb^All^). LHb^All^ neurons showed dynamic activity throughout the trial, and were significantly modulated by reward outcome as well as several other behavioral events (Figure 1H-K, Figure S1H-J), consistent with previous reports^15,16,19,20,49^.

The absence of LHb^Tac1^ modulation outside of reward omissions was striking, suggesting LHb^Tac1^ neurons might preferentially encode omissions in the task. To verify LHb^Tac1^ tuning to omissions, we used a series of behavioral assays in LHb^Tac1^ and LHb^All^ recordings to test whether (a) LHb^Tac1^ omission responses generalize across reward paradigms and (b) LHb^Tac1^ neurons are also recruited by innately aversive stimuli. First, we trained mice on a different reward-guided task consisting of a cued reward conditioning paradigm in which mice learn to associate two auditory cues with either high or low probability of reward (Figure 2A-B). During each trial, a 4 kHz or 13 kHz tone is pseudo-randomly presented for 1 s, then a reward is delivered or omitted at the reward port. One cue predicts likely reward (80% reward, 20% omission) and the other predicts likely omission (20% reward, 80% omission). To temporally segregate responses to cues and outcomes, cues were only delivered when the mouse was on the opposite side of the chamber. The pairing of auditory cue frequencies and reward contingencies was counterbalanced across mice, and there were no differences in the behavioral responses to the two frequencies (Figure S2A, E). All mice formed an association between the reward-predictive cues and their associated outcomes, displaying shorter latencies to approach the reward port following high probability cues (Figure 2C). As in our poke-reward task, we saw a significant increase in activity in LHb^Tac1^ neurons following reward omissions (Figure S2B), confirming that LHb^Tac1^ neurons encode reward omissions in both instrumental (poke-reward) and classical (cue-reward) conditioning.

**Figure 2.**
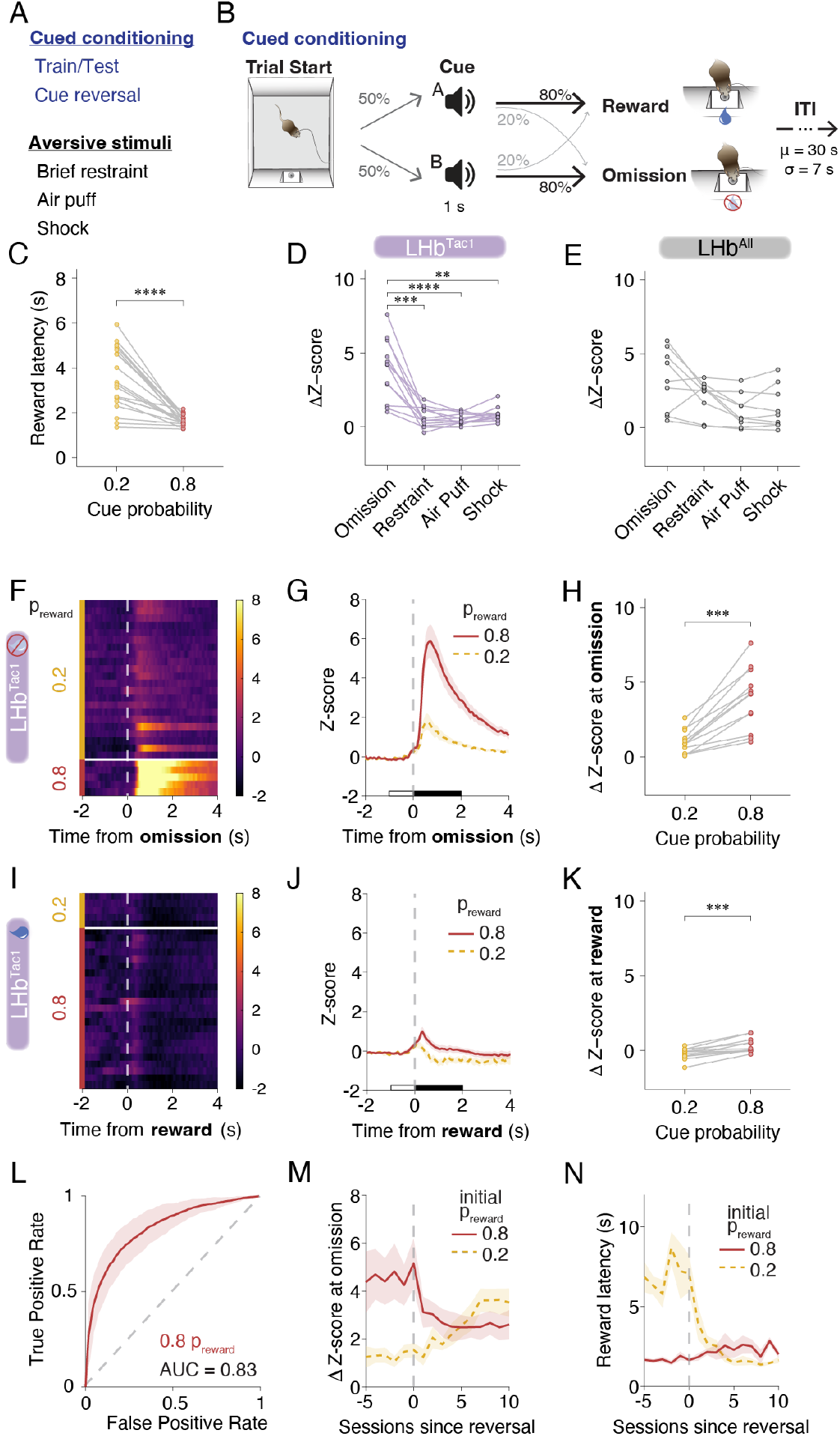
LHb^Tac1^ neurons preferentially signal reward outcomes in an expectation-dependent manner. (A) Behavioral assays for fiber photometry recording in reward and aversion. Handling-naïve animals first undergo brief restraint (30 s). Animals are water restricted for the auditory cued conditioning and then return to ad lib water and undergo aversive assays (air puff and mild foot shock). (B) Cued reward conditioning task. Mice are trained to associate an auditory tone (4 kHz or 13 kHz) with probabilistic sucrose water rewards. Cues predict either 80% or 20% probability reward. (C) Average latency from onset of auditory cue to reward port entry separated by cue-associated reward probability (paired t-test, p < 0.0001, n = 21). (D-E) Average responses to behavioral stimuli for LHb^Tac1^ (D) and LHb^All^ (E) mice, including reward omissions during cued conditioning, brief handling restraint, air puff, or foot shock. Responses calculated as mean z-score during stimulus minus mean z-score during 1s before stimulus onset, or 5 s before stimulus onset for restraint. Analysis windows for the stimuli were adjusted to reflect their duration: 1 s for shock and air puff; 2 s for reward omission; 5 s for restraint. LHb^Tac1^, n = 9; LHb^All^, n = 12. Details of multiple comparisons in Table S1). (F) Example of z-scored fluorescence of LHb^Tac1^ neurons aligned to reward omission for trials following either low (top) or high (bottom) probability cues. (G) Average z-scored fluorescence aligned to reward omission during 80% p_reward_ cue trials (solid red) and 20% p_reward_ cue trials (dashed yellow). (H) Average responses to reward omissions calculated as mean z-scored fluorescence 2 s following reward port entry minus 1 s preceding reward port entry (paired t-test, p < 0.001). (I) Example of z-scored fluorescence of LHb^Tac1^ neurons aligned to reward delivery for trials following either low (top) or high (bottom) reward probability cues. (J) Average z-scored fluorescence aligned to reward delivery during 80% p_reward_ cue trials (solid red) and 20% p_reward_ cue trials (dashed yellow). (K) Average responses to reward delivery calculated as mean z-scored fluorescence 2 s following reward port entry minus 1 s preceding reward port entry (paired t-test, p < 0.001). (L) Receiver operator characteristic (ROC) curve for a support vector machine trained to predict the identity of the reward-predictive cue using the time series of neural activity and the reward outcomes as predictors for each mouse. Curve indicates mean true positive and false positive rates for the high probability cue type across mice. AUC = 0.83. n = 12 animals (M) Average z-scored responses to reward omissions across sessions relative to reversal of cue-reward contingencies (n = 5 animals). (N) Average latency from cue onset to reward port entry separated by cue-associated reward probability across sessions relative to reversal of cue-reward pairing (n = 5 animals). Error bars denote SEM. For all panels, * p < 0.05, ** p < 0.01, *** p < 0.001, **** p < 0.0001

These data are consistent with the well-established role of LHb in reward-guided behavior^50,51^, but the habenula also demonstrates strong activation in response to innately aversive stimuli^30,52–54^ . To assess the selectivity of LHb^Tac1^ tuning to reward omission, the same mice were exposed to a series of aversive stimuli including temporary restraint, air puff, and mild inescapable shock. LHb^Tac1^ activity at reward omission in the cue-reward conditioning task was higher than at any of the aversive stimuli, which did not differ significantly from each other (Figure 2D, Table S1, S2). In contrast, LHb^All^ neurons were recruited by all stimuli, showing no consistent differences across reward omissions, restraint, air puff, and shock (Figure 2E, S2F, Table S1, S2). Together, these data suggest that LHb^Tac1^ neurons may represent a genetically defined subpopulation of LHb neurons preferentially tuned to reward omissions, rather than broadly encoding negative valence stimuli.

Considering the striking specificity of the LHb^Tac1^ population for encoding the omission of an expected reward in both reward tasks, we next tested whether Tac1 might represent a cell type marker for nRPE encoding neurons. If so, we would expect (a) that changes in both the predicted value and realized value would alter activity in LHb^Tac1^ neurons and (b) LHb^Tac1^ activity would scale with the discrepancy between the prediction and the outcome.

### Predictive cues modulate reward outcome responses in LHb^Tac1^ neurons

Predictions about the value of potential rewards can be shaped by sensory cues in the environment. In the cued conditioning task, the presentation of two cues associated with different reward probabilities changes the expected value of reward on a trial-by-trial basis. For example, when a cue predicts a high probability of reward, it generates a strong expectation for reward; if the reward is then omitted, the resulting prediction error is more negative. When separating the outcome responses by the cues that preceded them, we found significant differences in the neural response in the two trial types. Omission-evoked activity was elevated following high probability cues relative to low probability cues, and reward-evoked activity was suppressed following low probability cues relative to high probability cues (Figure 2F-K). Both features are consistent with RPE encoding. Using the activity at reward outcomes (delivered and omitted), we were able to classify the identity of the preceding cue with 75% and 77% accuracy for the low and high probability cues, respectively (Figure 2L). When the auditory cue-outcome pairings were reversed, omission-evoked activity reflected the updated cue-reward contingency, consistent with dynamic remapping of reward associations (Figure 2M). The reversal was incomplete (p = 0.05, n = 5), which may reflect a difference in the acquisition and extinction rate of the cue-outcome pairing, as suggested by the asymmetrical shift in behavioral responses to the two cues after reversal (Figure 2N). Together, the data indicate that LHb^Tac1^ represents a genetically-defined subpopulation of LHb neurons tuned to reward outcomes and modulated by reward expectation.

### Prior reward history modulates LHb^Tac1^ responses to reward omissions

Reward predictions are driven by both external cues, as demonstrated in the cued conditioning task, as well as internal estimations derived from recent reward history. To test if LHb^Tac1^ reward outcome responses were also sensitive to history-based predictions of reward value, we systematically varied the probability of reward between sessions in the poke-reward task (p_reward_ = 0.5, 0.7, or 0.9). We found that reward omissions in sessions with a higher reward probability, and thus a larger expected value, triggered stronger responses in LHb^Tac1^ neurons than omissions in sessions with low reward probability (Figure 3A-B). To understand the timescale over which animals update their predictions about the reward outcome, we examined trial-by-trial activity to quantify how recent reward outcomes affected omission responses within a session. We compared omission responses in trials preceded by rewarded delivery to those preceded by a reward omission. Activity during reward omissions was significantly lower when the previous trial was also omitted (Figure 3C-D), suggesting LHb^Tac1^ activity reflects underlying reward estimations that are updated on the timescale of individual trials. To determine how far back in time recent omissions influence LHb^Tac1^ activity in the ongoing trial, we used a linear model fit using the history of reward outcome (delivered, omitted) and performance (premature, correct, no response) as predictors. Recent omissions significantly reduce the LHb^Tac1^ activity at the current omission as far as 7 trials back, with the most recent rewardoutcomes having the strongest influence on the current trial (Figure 3E). In a similar reward-guided task, rewards were continuously delivered in a block structure rather than pseudo-randomly delivered throughout the session, revealing a cumulative effect of recent reward history where the reward omission response decreased with each subsequent omission in the block (Figure 3F-G). Across this same timeframe, the behavioral response rate decreased throughout the omission block as the mice disengaged from the task (Figure 3H). Together, these data suggest LHb^Tac1^ neurons are modulated not only by external predictive cues, but also by estimates of recent reward history and can integrate outcome information across trials.

**Figure 3.**
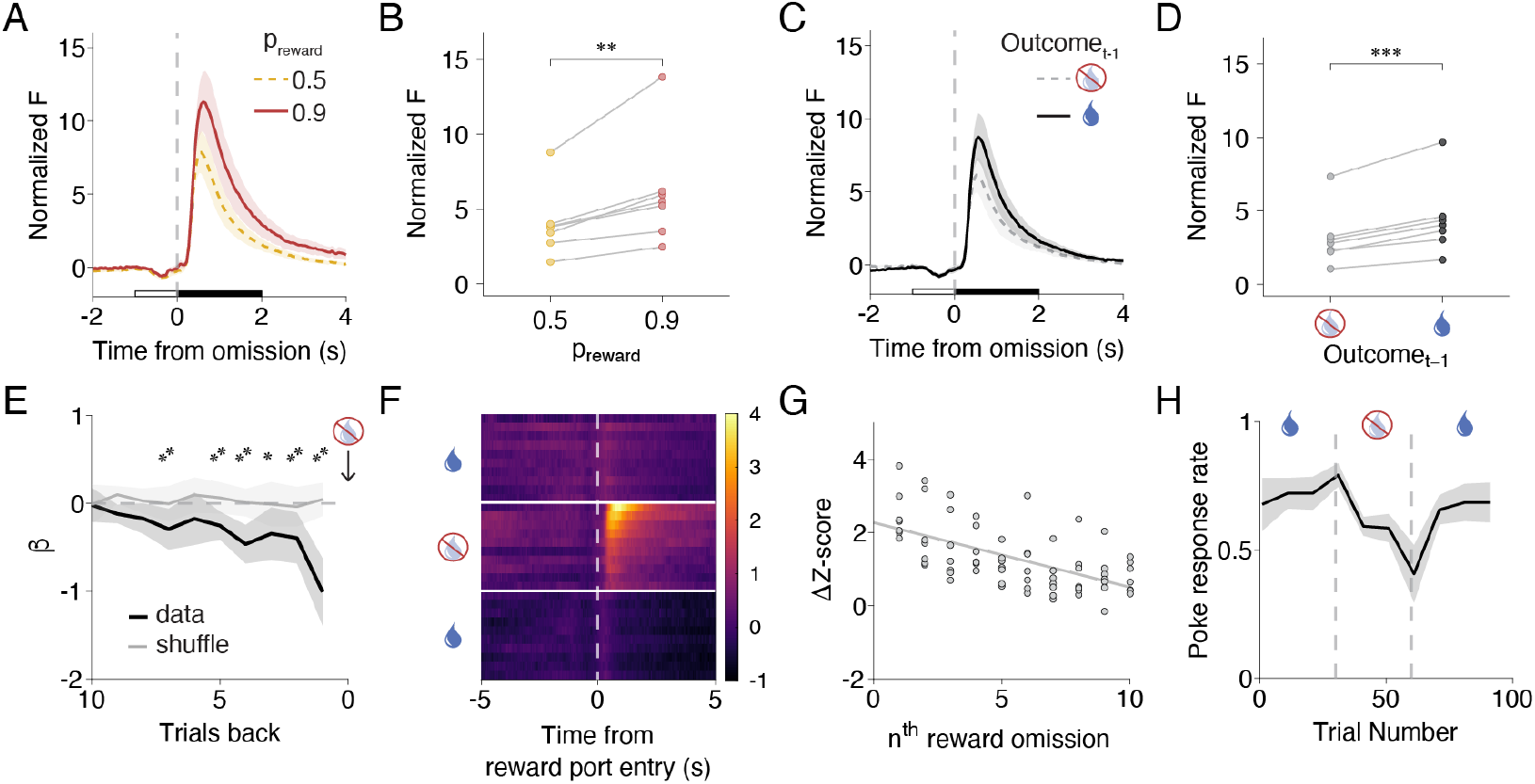
LHb^Tac1^ omission responses are modulated by reward history. (A) Averaged normalized fluorescence aligned to reward omissions during poke-reward task in sessions with reward probability 0.9 (solid red) and 0.5 (dashed yellow). (B) Average response to reward omissions separated by session reward probability calculated as mean fluorescence 2 s after reward port entry (filled bar) minus mean fluorescence 1 s before reward port entry (open bar). Paired t-test, p < 0.01. (C) Averaged response aligned to reward omissions during poke-reward task sessions with p_reward_ = 0.5, separated by reward outcome of previous trial. Black, outcome_t-1_ delivery; dotted grey, outcome_t-1_ omission. (D) Average response to reward omissions separated by previous trial outcome calculated as mean fluorescence 2 s after reward port entry minus mean fluorescence 1 s before reward port entry. Paired t-test, p < 0.001. (E) Average coefficient weights of a linear mixed effects model trained on the reward delivery on the past 10 trials in each animal. Black line indicates the median of across animals; gray line indicates the median of a model trained on shuffled reward history. p values derived from a permutation test of the difference in medians. Shaded error bars indicate standard deviation of coefficients across animals. (F) Variant of the poke-reward task with 3 nose poke ports and short (1 s) cue durations that contained a block of reward omissions on Trials 31-60. Fiber photometry data aligned to reward port entry during correct trials before, during and after the omission block, as indicated on the left. Color bar denotes the z-scored fluorescence. (G) Linear regression of the reward omission response across the omission block. R^2^ = 0.35, p < 0.0001. (H) Behavioral engagement during the omission block in the same animals as F-G, calculated at the number of no response trials, binned every 5 trials and averaged across animals, p < 0.01, R^2^ = .18 by linear regression. For all panels, n=7 and error bars indicate SEM, except in (E) where error bars indicate standard deviation.

### LHb^Tac1^ activity scales with the magnitude of nRPE

The experiments described above compare activity at reward omission under varying expectation. If LHb^Tac1^ activity reflects a prediction error signal, LHb^Tac1^ neurons should also respond when rewards are delivered, but are smaller-than-expected, with activity scaling proportionally to the deviation from the expected reward size. To test this prediction, we ran trained mice on a variant of the poke-reward task where reward was delivered on all correct trials, but the reward size was variable: 5 µL, 10 µL, and 30 µL rewards were delivered in 15%, 70%, and 15% of trials, respectively. In this paradigm, mice were well-trained to expect 10 µL prior to test sessions, and we interleaved all test sessions with 10 µL-only sessions to keep the expected value as stable as possible, leading to trials in which the computed RPE was negative (5 µL), zero (10 µL), or positive (30 µL). We found that LHb^Tac1^ activity scaled with nRPE: worse-than-expected outcomes (5µL) showed increased activity compared to neutral (10µL) and better-than-expected (30µL) outcomes (Figure 4A-B). A multiclass model trained on the activity at reward outcome was able to predict the reward size delivered above chance (AUC = 0.94, 0.74, and 0.85 for 5 µL, 10 µL, and 30 µL, respectively; Figure 4C).

**Figure 4.**
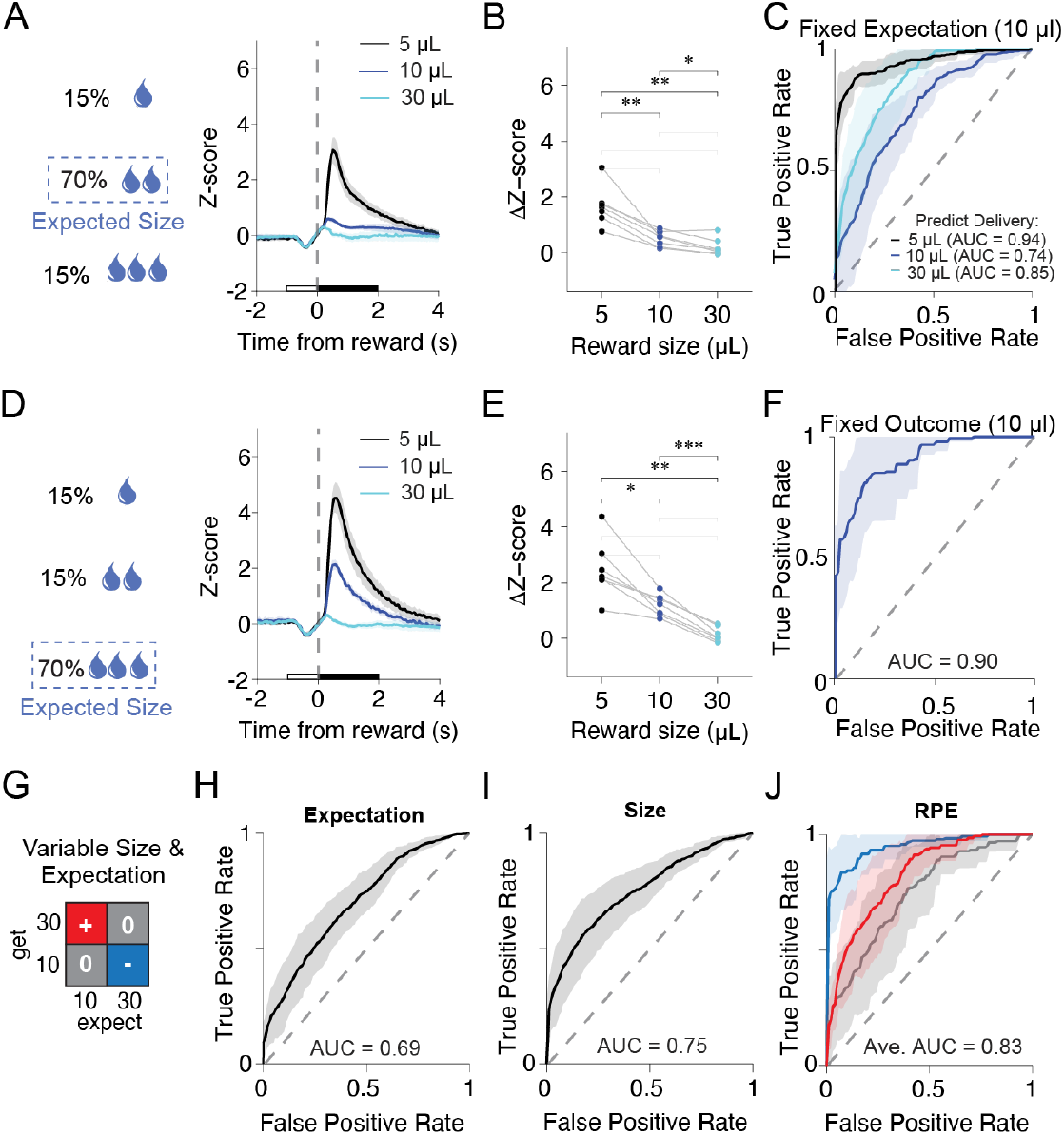
LHb^Tac1^ activity scales with the magnitude of prediction errors. (A-C) Poke-reward task variant where animals are trained to expect 10 µL rewards and then undergo probe sessions where variable 5 µL, 10 µL, and 30 µL rewards are delivered in 15%, 70%, and 15% of trials respectively. (D-E) Poke-reward task variant where animals are retrained to expect 30 µL rewards and then undergo probe sessions where variable 5 µL, 10 µL, and 30 µL rewards are delivered in 15%, 15%, and 70% of trials respectively. (A,D) Averaged activity aligned to reward delivery for 5 µL (black), 10 µL (dark blue) and 30 µL (cyan) rewards in sessions where animals either expect 10 µL (A) or 30 µL (D) rewards. n = 7 animals. Error bars indicate SEM. (B,E) Average reward-evoked activity in 5 µL, 10 µL, and 30 µL trials in sessions where animals either expect 10 µL (B) or 30 µL (E) rewards. Average calculated as the mean z-scored fluorescence 2 s after reward port entry minus 1 s before reward port entry. ANOVA, p < 0.01. Details of multiple comparisons in Table S3B-C. (C) ROC curves for a multiclass support vector machine trained to predict reward size using neural activity as a predictor in sessions where animals expect 10 µL rewards. 5 µL: AUC = 0.94; 10 µL: AUC = 0.74; and 30 µL: AUC = 0.85. n = 1232 trials across 28 sessions in 7 animals. (F) ROC curves for a support vector machine trained to predict expected size from the neural activity at 10 µL reward deliveries across both reward expectation sessions. AUC = 0.90. (G-J) ROC curve for a multiclass support vector machine trained to predict either the expected reward size (H), the delivered reward size (I), or the RPE associated with the expect::get pairing (J). Training data consisted of the timeseries of the neural response in LHb^Tac1^ neurons at reward outcomes for 10 µL and 30 µL reward deliveries across 10 µL and 30 µL expectation. RPE calculated as the delivered value minus the expected value, resulting in RPE of -20, 0, and 20; Average AUC = 0.92, 0.72, and 0.83, respectively. n = 2546, 2078, and 546 trials, across 7 mice. Blue, red, and grey denote nRPE, pRPE, and no RPE respectively. For all panels, * p < 0.05, ** p < 0.01, *** p < 0.001, **** p < 0.0001

Consistent with our results in the probabilistic poke-reward task (Figure 3E), we found the history of recent reward sizes affects ongoing trials, with significant trial history effects of reward size two trials back (p < 0.001, n = 7, Figure S3C). To more systematically probe whether LHb^Tac1^ activity was sensitive to changes in both reward size and expectation, in the same cohort of animals, we retrained mice over several sessions to expect 30 µL rewards and measured LHb^Tac1^ responses to probe trials with 5 µL and 10 µL rewards, producing nRPE of two different magnitudes. This allowed us to test two predictions. First, if LHb^Tac1^ neurons encode prediction errors, the magnitude of the neural response should scale with the difference between reward expectation and reward outcome. Second, we would predict that the response to a fixed outcome across two different expectations (expect 30 µL vs expect 10 µL) would differ. Consistent with nRPE encoding, we found significant effects of realized reward size, expected reward size, and the interaction between the two (Table S3A). LHb^Tac1^ activity scaled with nRPE, showing significantly more activity at 5 µL rewards than at 10 µL rewards (Figure 4D-E, Table S3B). Additionally, both 5 µL and 10 µL rewards elicited stronger responses in LHb^Tac1^ neurons when animals expected 30 µL rewards compared to 10 µL rewards (Figure 4A,D; Table S3C) and a classifier trained on 10 µL deliveries across the two experiments was able to predict the expected reward size (Figure 4F, AUC = 0.90). Attempts to simultaneously predict both expected and delivered reward size when both parameters varied across the data set yielded informative classification failures. We found that trials with the same RPE, e.g. *expect 10: get 10* vs *expect 30: get 30*, were routinely confused (Figures S3D), suggesting that a model predicting the computed RPE for a given trial may outperform models predicting the expected or realized reward size alone. To test this idea, we compared all three classifiers (size, expectation, and RPE) on a data set where both size and expectation varied (Figure 4G). The model trained to predict computed RPE on a given trial performed better than those predicting expectation or size alone (average AUC = 0.69, 0.75, and 0.83 for expectation, size, and RPE, respectively), demonstrating that the activity of LHb^Tac1^ neurons encodes RPE. Consistent with our data from both the variable size paradigm (Figure 4C) and the cued conditioning task (Figure 2J), model performance was better for negative RPE than positive RPE. Taken together, we find the activity of LHb^Tac1^ neurons encodes the difference between expected and realized reward value, showing that LHb^Tac1^ neurons preferentially and proportionally encode negative prediction errors.

## DISCUSSION

The functional diversity of the LHb is well established at the individual unit level, including encoding of reward prediction error^15–17,26,27^ and a variety of aversive stimuli^18,29,30,52,54,55^. However, few studies have linked the subpopulations mediating these behaviors to genetically-defined cell types^39,45,47,56^. Here, we measured population activity from tachykinin 1-expressing habenular neurons in mice during reward-guided tasks. We found that LHb^Tac1^ neurons consistently encode reward outcomes across multiple reward-learning contexts, with outcome responses scaling with the magnitude of nRPE. LHb^Tac1^ neurons exhibited only weak responses to innately aversive stimuli that are known to engage a broader set of LHb neurons. Together, these data demonstrate that LHb^Tac1^ neurons represent a genetically-defined subpopulation of habenular neurons that preferentially encodes nRPE.

### Valence-biased RPE encoding

Traditional RL theory assumes reward prediction error is represented on a symmetric, linear scale centered around a single quantity^57^ . However, neural recordings from mice, nonhuman primates, and humans reveal substantial diversity in RPE encoding across neurons in the VTA^8^, PFC^58^, ACC^59,60^, and LHb^15,16,18^ . We found that LHb^Tac1^ neurons encoded negative RPEs more reliably than positive RPEs across multiple reward contexts. The high-reward context of our assays, all of which were at or above a 50% reward rate, may have limited our ability to detect changes at positive RPEs. Alternatively, the asymmetry may reflect biological rather than experimental factors. Although single unit recordings of LHb^Tac1^ cells are limited, they suggest a median firing rate of ∼4 Hz^61^, raising the possibility that weak pRPE encoding could be driven by the biophysical constraints of a floor effect. LHb lesions impair negative RPE signaling in DA neurons but left positive RPEs intact^27^, in line with our finding of asymmetric encoding in LHb^Tac1^ neurons and in support of a model where LHb^Tac1^ neurons may preferentially contribute to nRPE-induced suppression of DA neuron firing. Recent computational advances suggest asymmetric RPEs provide a potential mechanism underlying differential learning rates associated with positive and negative outcomes, improving adaptation to contexts with variable reward availability^58,60,62–64^ .

### Peptidergic Modulation

LHb axons project to several midbrain and hindbrain regions^65^, including the VTA^66,67^, rostromedial tegmental nucleus (RMTg)^24,68^, and dorsal raphe^69^, positioning them to influence both dopaminergic and serotonergic signaling. LHb axons projecting to the midbrain predominately synapse on GABAergic neurons in the VTA and RMTg, which inhibit VTA dopamine neurons and shape their RPE signaling^24–26,66,67,70^. The cell-type specific projections of LHb^Tac1^ neurons have not been described, but their ability to encode nRPE suggests they may contribute to this disynaptic inhibitory LHb-RMTg-VTA pathway. Alternatively, they may route nRPE information to other known LHb targets, such as the dorsal raphe, which has been shown to modulate behavioral flexibility in reward guided paradigms, making it a compelling candidate target for error-encoding LHb^Tac1^ neurons^71–74^.

*Tac1* encodes the propeptide for Substance P and Neurokinin A, and their cognate neurokinin receptors are present in the LHb as well as its target regions (VTA and DR)^75–77^. It is unclear if activity at prediction errors is sufficient to trigger neuropeptide release from LHb^Tac1^ neurons^78^, leaving open questions for the role of peptidergic modulation in RPE encoding. Local somatodendritic release has not been studied extensively, although antagonist infusions into the habenula have been shown to reduce depression-like phenotypes^79^. Habenular lesions reduce Substance P levels in target regions, including the DR, suggesting axonal release may also occur^80,81^. Future work is needed to clarify how nRPE signals are routed to downstream regions and the extent to which peptidergic signaling contributes to these circuits to guide decision-making in dynamic environments.

### Implications for reward-guided decision-making

Growing evidence suggests the LHb plays a broader role in coordinating motivation and movement systems to drive multiple learning processes^54,82^, particularly in complex environments^83–85^. Associative learning requires binding predictive cues and outcomes. LHb^Tac1^ neurons reliably encoded cue-reward associations, with omission-evoked activity significantly modulated by cue identity. In situations where external cues do not guide decisions, animals must rely on internal representations of the environment’s reward state derived from past outcome history. We found LHb^Tac1^ neurons integrated outcome history across multiple trials. This suggests LHb^Tac1^ neurons could facilitate reward-guided decision-making even in environments with dynamic reward contingencies. Lesion or pharmacological inactivation of the LHb is sufficient to disrupt choice preferences during probabilistic reversal learning tasks in which animals must recognize changing reward contingencies and act accordingly^86^. However, these manipulations modulate all LHb signaling, not just nRPE-encoding neurons. By identifying Tac1 as a molecular marker for RPE-encoding neurons, our work provides a genetic handle for future studies to dissect this specific computation. Genetic approaches that harness Tac1 as a cell-type marker for nRPE neurons will serve as valuable tools for understanding how nRPE signals in the LHb contribute to learning in dynamic environments.

## ACKNOWLEDGEMENTS

We thank the entire Sylwestrak Laboratory for helpful discussions, Dr. Cristopher Niell and Dr. Matthew Smear for insightful feedback on the manuscript, and Dr. James Murray and Dr. Christian Schmid for advice on analysis. We also thank the University of Oregon Terrestrial Animal Care Services staff for animal care assistance, Adam Fries for imaging technical support, and the Stanford Vector Core for providing viruses. This work was funded by the Brain Behavior & Research Foundation NARSAD Young Investigator Grant (27735), the National Institute of Mental Health (Grant Nos. F31MH135679 to K.E.S. and RF1NS132914 to E.L.S.), and the Esther A & Joseph Klingenstein-Simons Fellowship Award.

## AUTHOR CONTRIBUTIONS

K.E.S. contributed to conceptualizing experiments, data curation, formal analysis, funding acquisition, project administration, software, validation, and writing the original manuscript draft, and conducted behavioral training and photometry recordings for operant and classical conditioning, probabilistic and variable reward experiments, innate aversion experiments, and Tac1 expression labeling. E.L.S. contributed to supervision, conceptualizing experiments, data curation, formal analysis, funding acquisition, project administration, resources, software, validation, and writing the original manuscript draft. T.A.S. contributed to classical conditioning and innate aversion experiments. B.H. contributed to probabilistic reward experiments. J.R.K. contributed to operant conditioning experiments. All authors contributed to review and editing of the manuscript.

## SUPPLEMENTAL INFORMATION

**Figure S1.**
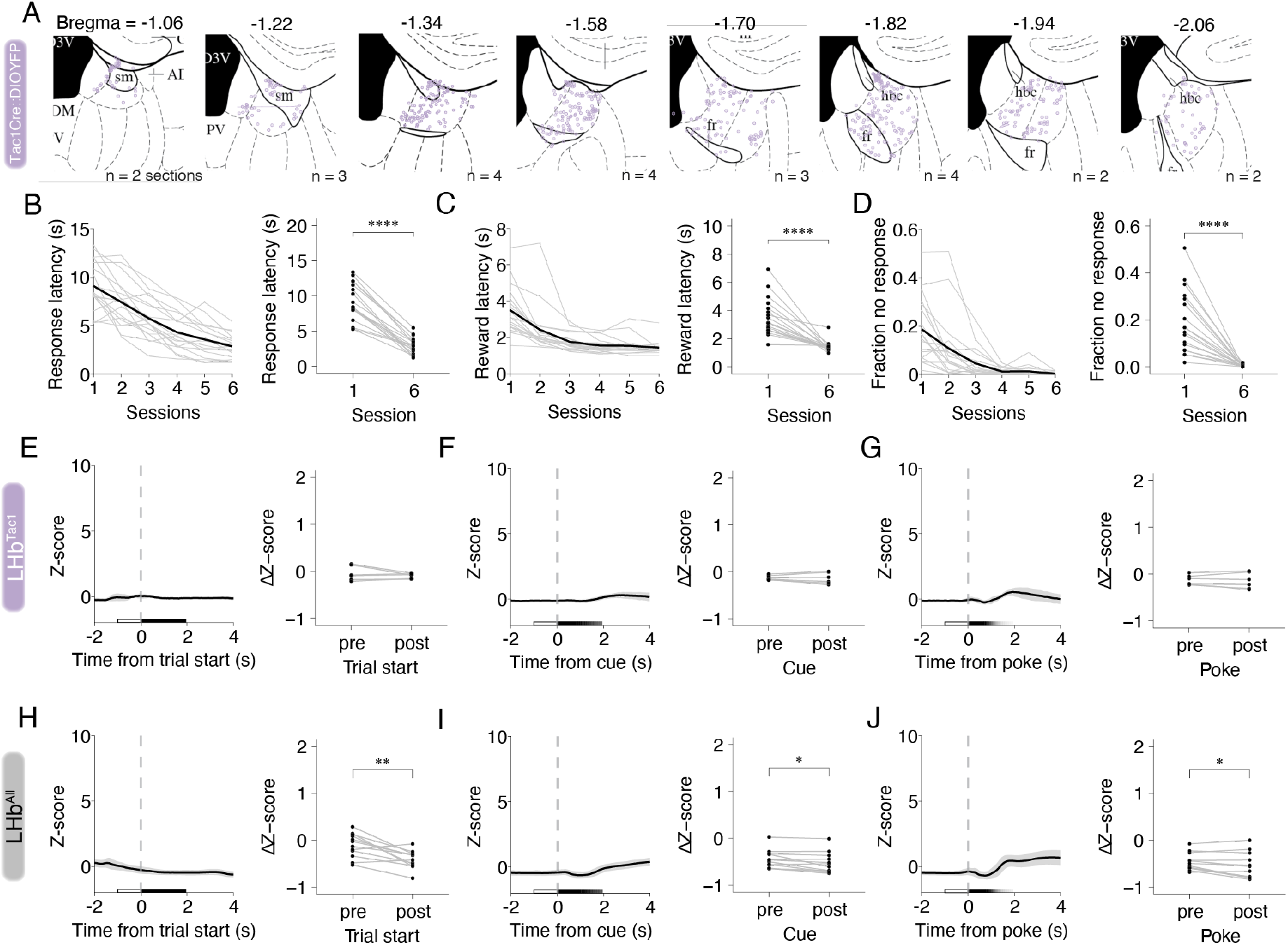
LHb^Tac1^ neurons preferentially respond to reward omissions. (A) Distribution of virally labeled *Tac1*^*+*^ neurons in Tac1-Cre mice injected with AAV-DIO-YFP. Sections were registered to the nearest atlas plate using n = 24 sections from 4 mice. (B-D) Performance metrics across training in the poke-reward task. Grey lines indicate session averages for each animal. Black lines indicate average across cohort (n = 18). Left panels, change in each behavioral metric over the first 6 sessions of training. Right panels, difference in each metric between session 1 and session 6. Response latencies (B) are defined as the time from cue onset to nose poke into the cue port (paired t-test, p < 0.0001). Reward latencies (C) are defined as the time from nose poke into the cue port to entry into the reward port (paired t-test, p < 0.0001). Fraction of “no response” trials (D) is defined as the proportion of trials in a session in which animals failed to poke in the cue port for the duration of the 30 s cue (paired t-test, p < 0.0001). (E-J) Average response of LHb^Tac1^ (E-G) or LHb^All^ populations (H-J) to different behavioral events in the poke-reward task. Left panels, times series data for fiber photometry signal at trial start (E,H), cue onset (F,I), and nosepoke into the cue port (G,J). Right panels, event evoked changes in activity, calculated as the difference in z-score between a baseline period (open boxes) and the analysis window (filled boxes). Paired t-test with FDR correction for multiple comparisons. Analysis window for each epoc (start, cue, poke) includes 2 s from event onset or time from behavioral epoc to entry into the reward port, whichever comes first. Gradient shaded boxes indicate average distribution of analysis windows across animals. For all panels, * p < 0.05, ** p < 0.01, *** p< 0.001, **** p < 0.0001.

**Figure S2.**
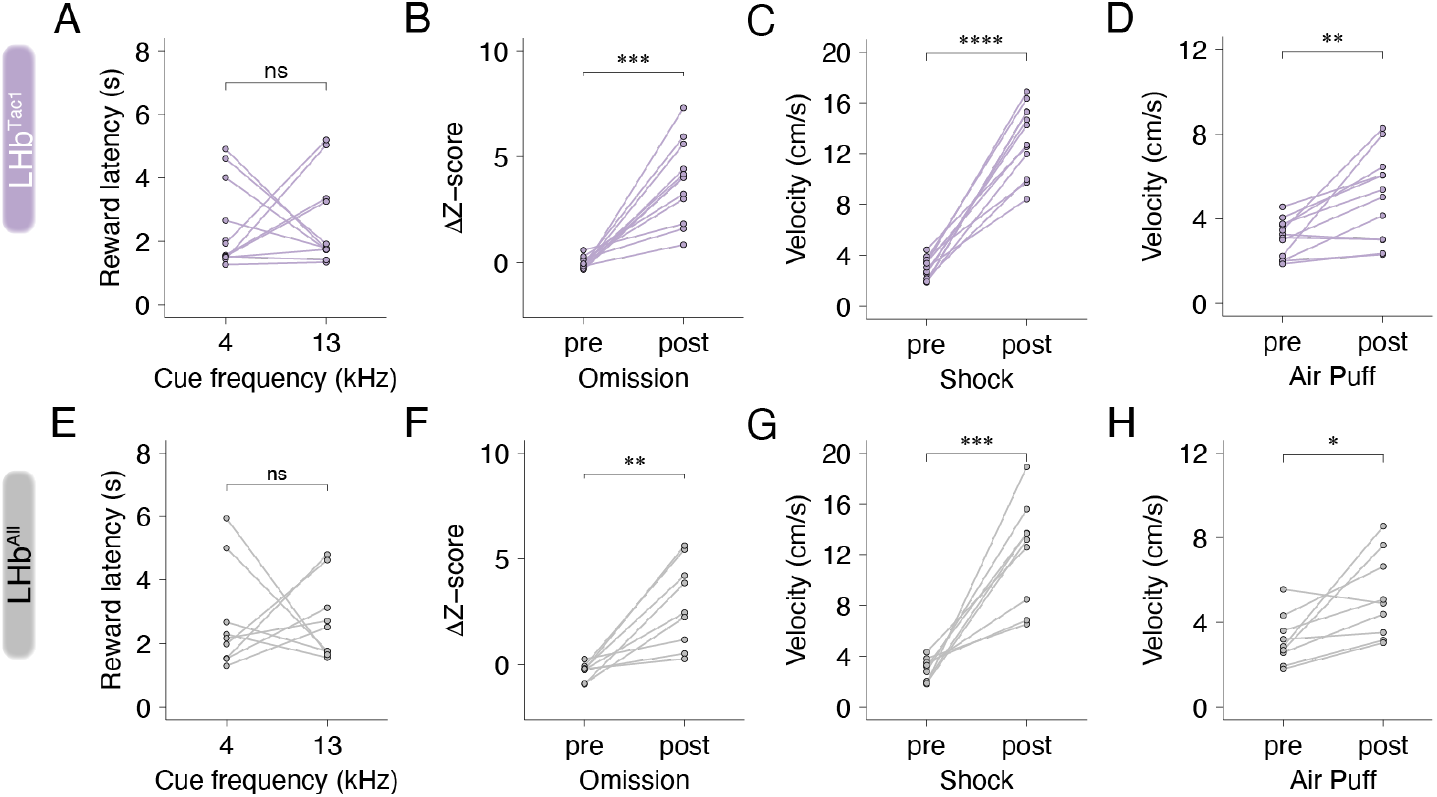
LHb^Tac1^ neurons preferentially signal reward outcomes in an expectation-dependent manner. (A,E) Average latency to reward port entry separated by auditory cue frequency in LHb^Tac1^ (A) and LHb^All^ (E) mice. (B, F) Change in activity at reward omissions following high probability cue in cue-reward conditioning task in LHb^Tac1^ (B) and LHb^All^ (F) mice. Response calculated as mean fluorescence 2 s after reward port entry minus mean fluorescence during 1 s before reward port entry. Paired t-test with FDR correction for multiple comparisons across paradigms. Details of multiple comparisons in Table S2. (C, G) Average velocity of animals 1 s before and 1 s after onset of foot shock. LHb^Tac1^, paired t-test p < 0.0001; LHb^All^, paired t-test p < 0.001. (D, H) Average velocity of animals 1 s before and 1 s after onset of air puff. LHb^Tac1^, paired t-test p < 0.01; LHb^All^, paired t-test p < 0.01. For all panels, * p < 0.05, ** p < 0.01, *** p< 0.001, **** p < 0.0001. LHb^Tac1^, n = 12; LHb^All^, n = 9

**Figure S3.**
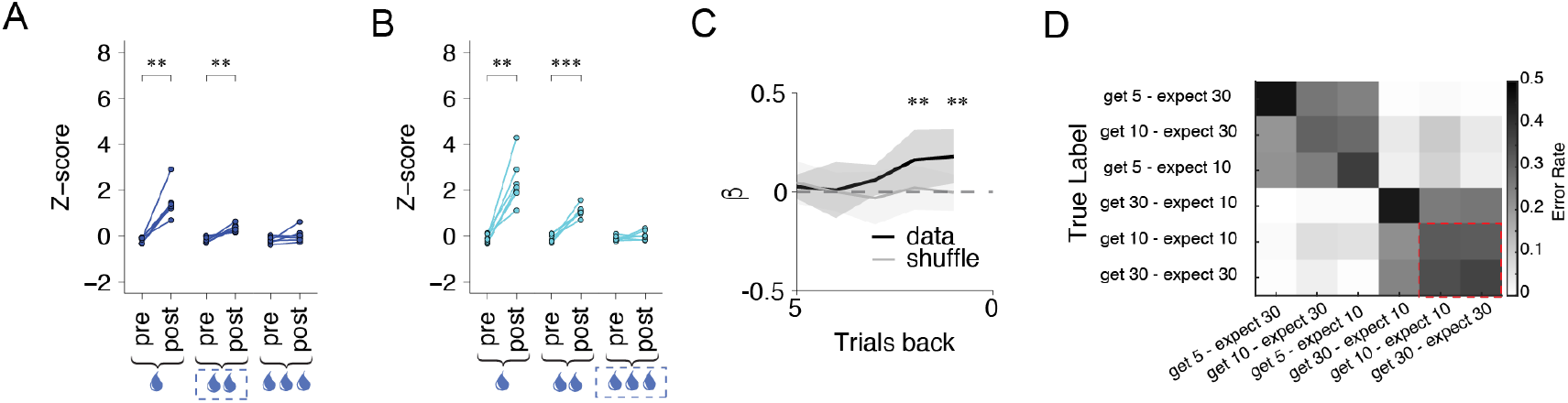
LHb^Tac1^ activity encodes RPE. (A-B) Average fluorescence before and after reward consumption during 5µL (left), 10 µL (middle) and 30 µL (right) rewards for each animal when expected size is 10 µL (A) or 30 µL (B), calculated in a 1 second window before the reward onset and 2 seconds after the reward onset, respectively. Paired t-test with bonferroni correction for multiple comparisons. 10 µL expected: 5 µL, p < 0.01, 10 µL, p < 0.01, 30 µL, p = 0.43; 30 µL expected: 5 µL, p < 0.01, 10 µL, p < 0.001, 30 µL, p = 0.63. Dotted blue square indicates expected reward size. (C) Trial history regression of reward size. Average coefficient weights of a linear mixed effects model trained on the reward size on the past 5 trials in each animal. Black line indicates the median across animals; gray line indicates the median of a model trained on shuffled reward history. p values derived from a permutation test of the difference in medians. Shaded error bars indicate standard deviation of coefficients across animals, n = 7. (D) Confusion matrix for joint classification of reward expectation and reward size with the time series of neural activity at reward delivery as a predictor. Grayscale indicates the fraction of each true label with that predicted label. Dotted red box indicates confusion between trial types with zero prediction error. Color mapping indicates the proportion of trials of a given expectation::outcome pairing with each predicted label.

**Table S1.** Statistical analysis of changes in fiber photometry fluorescence during reward omissions and innately aversive stimuli.

|  | Paradigm Comparison | Statistic | Adj. p-value | Significance |
| --- | --- | --- | --- | --- |
| **Tac1** | Omission, Restraint | -3.85 | $p < 0.001$ | *** |
| | Omission, Air Puff | -4.40 | $p < 0.0001$ | **** |
| | Omission, Shock | -3.41 | $p < 0.01$ | ** |
| | Restraint, Air Puff | $-5.54 \times 10^{-1}$ | $p = 1.00$ | ns |
| | Restraint, Shock | $4.37 \times 10^{-1}$ | $p = 1.00$ | ns |
| | Air Puff, Shock | $9.91 \times 10^{-1}$ | $p = 1.00$ | ns |
| **Pan-neuronal** | Omission, Restraint | -1.10 | $p = 1.00$ | ns |
| | Omission, Air Puff | -2.39 | $p = 0.10$ | ns |
| | Omission, Shock | -2.19 | $p = 0.17$ | ns |
| | Restraint, Air Puff | -1.30 | $p = 1.00$ | ns |
| | Restraint, Shock | -1.10 | $p = 1.00$ | ns |
| | Air Puff, Shock | $2.01 \times 10^{-1}$ | $p = 1.00$ | ns |

Average response to footshock and air puff calculated as average fluorescence 1 s after stimulus onset minus average fluorescence 1 s before stimulus onset. Average response to reward omissions calculated as average fluorescence 2 s after entry into reward port minus average fluorescence 1 s before entry into reward port. Average response to restraint calculated as average fluorescence 5 s after restraint onset minus average fluorescence 5 s before restraint onset.
Fluorescence z-scored across session. Data represents average across animals.

**Table S2.** Statistical analysis of changes in fiber photometry fluorescence at reward omissions and innately aversive stimuli.

|  | Paradigm | Statistic | Adj. p-value | Significance |
| --- | --- | --- | --- | --- |
| **Tac1** | Omission | -6.52 | $p < 0.001$ | *** |
| | Restraint | -3.47 | $p < 0.01$ | ** |
| | Air Puff | -4.96 | $p < 0.001$ | *** |
| | Shock | -5.72 | $p < 0.001$ | *** |
| **Pan-neuronal** | Omission | -4.53 | $p < 0.01$ | ** |
| | Restraint | -5.18 | $p < 0.01$ | ** |
| | Air Puff | -2.99 | $p < 0.05$ | * |
| | Shock | -2.73 | $p < 0.05$ | * |

Average fluorescence before and after reward omission, restraint, air puff, and foot shock. Responses calculated as mean fluorescence during stimulus minus mean fluorescence during 1s before stimulus onset, or 5 s before stimulus onset for restraint. Analysis windows for the stimuli were adjusted to reflect their duration: 1 s for shock and air puff; 2 s for reward omission; 5 s for restraint.
Fluorescence z-scored across session. Data represents average across animals.

**Table S3A.** Statistical analysis of changes in fiber photometry fluorescence during reward consumption: 2-Way Repeated Measures ANOVA.

| Effect | (DFn, DFd)* | F | GES | Adj. p-value | Significance |
| --- | --- | --- | --- | --- | --- |
| Reward size | 1.05, 6.3 | $3.11 \times 10^1$ | $6.89 \times 10^{-1}$ | $p < 0.01$ | ** |
| Expectation | 1, 6 | $2.85 \times 10^1$ | $1.75 \times 10^{-1}$ | $p < 0.01$ | ** |
| Reward size * Expectation | 1.26, 7.56 | $2.15 \times 10^1$ | $1.18 \times 10^{-1}$ | $p < 0.01$ | ** |

Average response to reward calculated as average fluorescence 2 s after reward port entry minus average fluorescence 1 s before reward port entry.
Fluorescence z-scored across session. Data represents average across animals.
\*Degrees of freedom adjusted using Greenhouse-Geisser correction for sphericity violations.

**Table S4B.**
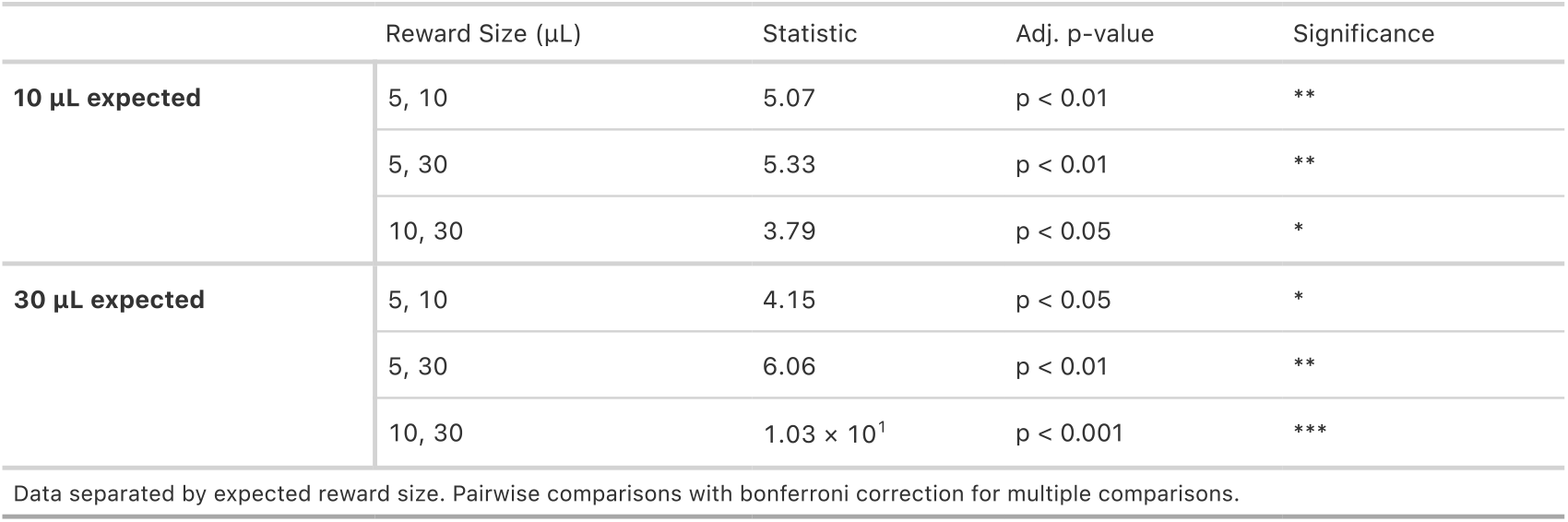
Post-hoc comparisons for delivered reward size effect.

**Table S3C.**
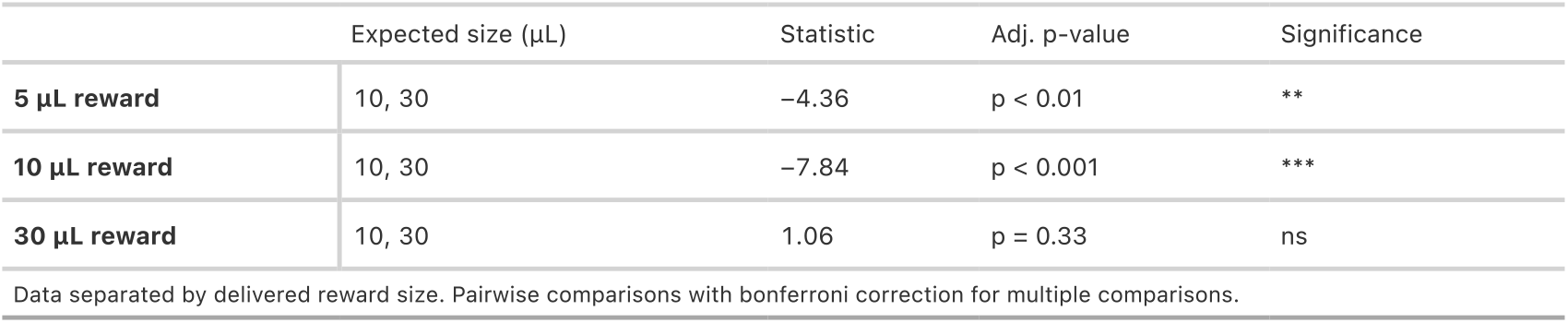
Post-hoc comparisons for expected reward size effect.

| | Expected size (μL) | Statistic | Adj. p-value | Significance |
| --- | --- | --- | --- | --- |
| **5 μL reward** | 10, 30 | -4.36 | $p < 0.01$ | ** |
| **10 μL reward** | 10, 30 | -7.84 | $p < 0.001$ | *** |
| **30 μL reward** | 10, 30 | 1.06 | $p = 0.33$ | ns |

Data separated by delivered reward size. Pairwise comparisons with bonferroni correction for multiple comparisons.

## STAR Methods

### Method Details

| Reagent/Resource | Source | Identifier |
| --- | --- | --- |
| **Antibodies** |  |  |
| Anti-GFP Polyclonal Antibody, Alexa Fluor 647 | Thermo Fisher Scientific | Cat # A-31852; RRID AB_162553 |
| Anti-GFP Polyclonal Antibody, Alexa Fluor 488 | Thermo Fisher Scientific | Cat # A-21311; RRID AB_221477 |
| **Bacterial and virus strains** |  |  |
| AAV8-Ef1a-DIO GCaMP 6f | Stanford Viral Vector Core | custom prep |
| AAV1-CAG-FLEX-jGCaMP8s-WRPE | Addgene (GENIE Project) | Addgene viral prep #162380-AAV1 |
| AAV1-Syn-GCaMP6m-WPRE-SV40 | Addgene (Douglas Kim and GENIE Project) | Addgene viral prep #100841-AAV1 |
| AAV1-CAG-Flex-GCaMP6m-WPRE-SV40 | Addgene (Douglas Kim and GENIE Project) | Addgene viral prep #100839-AAV1 |
| AAV1-Ef1a-DIO-EYFP | Addgene (from Karl Deisseroth) | Addgene viral prep #27056-AAV1 |
| **Chemicals** |  |  |
| 4% Perfusion Fixative | Thermo Fisher Scientific | Cat # 50-334-41 |
| PBS | Gibco | Cat # 70011044 |
| Triton X-100 | Sigma-Aldrich | Product # X100; CAS # 9036-19-5 |
| DAPI | Thermo Fisher Scientific | Cat # 00-4959 |
| Sucrose | Sigma Aldrich | Cat # S0389 |
| **Experimental models: Organisms and Strains** |  |  |
| B<sub>6</sub>; 129S-Tac1<sup>tm1.1(cre)Hze</sup> | The Jackson Laboratory | Strain # 021877; RRID: IMSR_JAX:021877 |
| C57BL/6 | The Jackson Laboratory | Strain # 000664; RRID: IMSR_JAX:000664 |
| **Software and algorithms** |  |  |
| Synapse | Tucker-Davis Technologies | https://www.tdt.com/component/synapse-software/ |
| FIJI | Open Source | https://imagej.net/software/fiji/ |
| MATLAB | MathWorks | https://www.mathworks.com/products/matlab.html |
| SLEAP<sup>87</sup> | Pereira et al., Nature Methods, 2022 | https://github.com/talmolab/sleap |
| BehaviorDEPOT<sup>88</sup> | Gabriel et al., eLife, 2022 | https://github.com/DeNardoLab/BehaviorDEPOT |

### Animal Behavior

#### Subjects

All experiments, surgical procedures, and animal care and euthanasia were in performed in accordance with the University of Oregon Animal Care and Use Committee guidelines and the National Institutes of Health’s Guide for the care and use of laboratory animals. All experiments were performed using adult mice (11-18 weeks). This study includes both male and female mice. For fiber photometry experiments, Tac1-Cre (B6; 129S-Tac1tm1.1(cre)Hze) and C57BL/6 mice were used.

#### Housing and handling

Prior to experiments, animals were housed in a temperature-controlled vivarium under scheduled lighting conditions (12 h light/dark cycle) and provided unrestricted access to food and water. In preparation for behavior testing, animals were transferred to reverse light cycle rooms at least 1 week prior to testing. Animals were handled daily for at least 3 days prior to the onset of behavior testing. All behavioral experiments were performed ZT14 – ZT20.

#### Water Restriction

Mice were weighed for three consecutive days prior to the onset of water restriction and weight was monitored daily following the onset of water restriction to ensure mice maintained >80% of their body weight. Mice were acclimated to sucrose by allowing animals already under water restriction temporary access to 10% sucrose solution until animals consumed their daily allotment of water.

### Operant behavior

#### Behavior chamber

Operant behavior experiments were conducted in custom-built behavior chambers (15 cm x 15 cm x 30 cm) equipped with behavioral ports on opposing ends of the chamber with a photogate, white LED, and solenoid valve for 10% sucrose water delivery. Chambers were placed inside sound attenuating boxes equipped with an LED house light and an overhead camera for behavioral monitoring. Tasks were controlled via an open-source, MATLAB-based data acquisition system and microcontroller (Bpod State Machine r2, Sanworks, https://github.com/sanworks/Bpod). The state machine recorded behavioral responses across trials and controlled the state of the task including LEDs, reward delivery, and house lights according to the task protocol. *Acclimation*. Animals were acclimated to the operant box for 10 min/day for 3 consecutive days during which time the house lights were illuminated. *Magazine*. On day 4, mice underwent magazine training during which 20 µL sucrose water rewards were available every 30 s in an illuminated reward nose poke port until animals consumed 50 rewards. House lights were illuminated throughout. *Operant training*. When animals retrieved water rewards with low latency (< 5 s) in the magazine stage, animals moved to operant training. After an ITI period (µ= 4s; σ = 0.2s), animals were required to poke into an illuminated nose poke port (initiation port) on the opposing side of the box, which resulted in illumination reward port light and delivery of a 10 µL reward. Premature pokes in the initiation port resulted in a 10 s timeout period during which house lights were extinguished. Latencies to retrieve reward longer than 120 s resulted in the start of a new trial (<0.5% of trials in trained animals). For most sessions, animals performed 115 trials, or 60 min whichever came first. In sessions with poor performance additional trials were run to enable animals to reach their daily quota for water to maintain weight. If additional water was required to maintain weight the remaining water was delivered >1 h after the end of the training session.

For experiments with a block of omitted rewards (Figure 3F), a modified operant task was used to increase sensitivity to changes in task engagement. In this task, 3 initiation ports are active instead of one. In addition, the LED cue duration is short (1 s), increasing attentional demands in the task. Behavioral shaping and training was performed as described previously^61^. Briefly, after magazine training, animals proceeded to a nonspatial session where all three initiation port LEDs were illuminated. After criterion, they advanced to spatial training, in which only one cue port was active for each trial, starting with a 30 s cue duration and progressively shortening the cue to 1 s. Training sessions consisted of 100% reward delivery. In test sessions with blocks of omitted trials, Trials 1-30 were 100% delivered, trials 31-60 were 0% rewarded, and trials 61-100 were 100% rewarded. For experiments manipulating session reward probability or size, the order of the sessions was counterbalanced across animals and test sessions were interleaved with training sessions with either 100% reward delivery or 10 µl fixed reward size.

### Auditory cue-reward conditioning

#### Behavioral chamber

Auditory cue-reward conditioning experiments were conducted in a similar chamber as used for operant behavior but with no initiation port. Auditory cues were generated using an Arduino-based device (Bpod HiFi module, Sanworks) and presented through two speakers inside the sound attenuating box. *Acclimation*. Animals were acclimated to the conditioning chamber for 10min/day for 3 consecutive days during which house lights were illuminated. *Magazine*. On Day 4, 20 µL sucrose water rewards were available every 30 s in the reward port until animals consumed 50 rewards. A 70 dB white noise tone was delivered for 1 s concurrent with head entry in the reward port. After they reached criterion (50 rewards in 30 minutes), mice were moved to cue-reward association training. *Auditory cue-reward association*. Following completion of magazine training, mice were trained to associate white noise presentation with sucrose water delivery at the reward port. To temporally separate the cue and reward and discourage camping near the reward port or premature licking prior to cue onset, animals were required to cross an IR photobeam located near the back of the box to initiate a trial. Crossing the photobeam triggered presentation of white noise for 1 s at 70 dB concurrent with delivery of 10% sucrose water reward in the reward port, after which animals retrieved reward. Trials ended with a variable inter-trial interval (µ = 30 s, σ = 7 s). Animals completed two sessions of cue-reward conditioning before advancing to probabilistic cue-reward conditioning. *Probabilistic cue-reward conditioning*. Probabilistic training consisted of two 70 dB pure tones of 4 kHz and 13 kHz where one cue predicts likely reward (80% reward, 20% omission) and the other cue predicts likely omission (20% reward, 80% omission). The tone frequency and associated reward probability were counterbalanced across mice to ensure that differences in behavioral responses and neural activity were independent of the tone frequency. Trials for the two cue types were pseudo-randomly interleaved in a 1:1 ratio. At the start of each trial, an animal crossing the IR photobeam triggered presentation of a 70 dB pure tone at either 4 kHz or 13 kHz for 1 s. Concurrent with the auditory cue, a 20 µL sucrose water reward was delivered in the reward port on rewarded trials after which animals retrieved reward. During unrewarded trials, a sham solenoid valve opened to mimic the sound of the solenoid valve. Latencies to enter the reward port longer than 30 s resulted in automatic transitions to the ITI. Trials ended with a variable inter-trial interval (µ = 30 s, σ = 7 s). For most sessions animals performed 60 trials or 60 min, whichever came first. If additional water was required to maintain weight the remaining water was delivered > 1 hr after the end of the training session. Animals completed 15 sessions of probabilistic training before cue-reward contingencies were reversed.

### Temporary Restraint

Animals were placed in an open field chamber and allowed to explore freely for 1 minute. Animals were then manually restrained for 30 seconds before being returned to the open field chamber for 30 seconds.

### Air Puff

Animals were placed in a chamber (18 cm x 18 cm x 30 cm) with a metal grated floor and allowed to explore freely throughout the session. An air diffuser box positioned below the chamber floor was connected to house air. Air puffs covering the full field of the chamber were delivered at 60 psi for 1 s using a pneumatic valve controlled by a Bpod behavioral control system. Trials were separated by a variable inter-trial interval (µ = 30 s, σ = 5 s). Air puff trials and sham trials in which a dummy valve was opened were presented with equal likelihood and pseudo-randomly interleaved in the session. Animals were exposed to 20 air puffs during a single session.

### Foot shock

#### Behavioral chamber

Animals were placed in a chamber (18 cm x 18 cm x 30 cm) with a metal grated floor. Foot shocks were delivered to the grid floor by a precision animal shocker (Lafayette Instrument, Model HSCK100AP). Foot shocks and speakers were controlled by an Arduino-based behavioral control protocol (BehaviorDEPOT^88^). LED strips were fixed to the top of the external chamber to illuminate the behavioral chamber for video capture. *Pre-exposure*. On the day prior to fear conditioning, animals were placed in the behavioral chamber and allowed to freely explore for 10 minutes without the presence of any shocks. *Foot shock*. Sessions started with a 30 s habituation period preceding the first trial. For each trial, a mild foot shock (0.1 mA) was administered for 1 s. Each shock presentation was followed by a variable inter-trial interval or 60 – 120 s. Animals were exposed to a single session with a total of 20 shocks.

### Fiber Photometry

#### Surgical Procedures

Male and female mice age 11-18 weeks were anesthetized with 4% isoflurane and maintained at 1-2% isoflurane mixed with oxygen at a flow rate of 1.5 – 2 L/min. Depth of anesthesia was monitored throughout surgery using tail pinch responses and breathing rates. Mice were mounted on a stereotaxic surgical station (David Kopf Instruments). 0.5 mL of saline and 1 mg/kg of Buprenorphine SR (Wedgewood Pharmacy) or 3.25 mg/kg Buprenorphine extended release (Ethiqa XR) were injected subcutaneously. A small craniotomy was made over the lateral habenula and stereotaxic injections were used to deliver viruses. For fiber photometry experiments, 200 – 300 nL of AAV-Ef1a-DIO-GCaMP6 or AAV-Syn-GCaMP6 was injected in the right or left LHb (AP: -1.35 mm; ML: +/-0.55 mm; DV: -0.315 mm) at a rate of 30 nL/min and a fiber optic cannula (Doric Lenses, MFC_400/430-0.66_3.5mm_MF2.5_A30) was placed above the lateral habenula (AP: -1.35 mm, ML: +/-0.55 mm, DV: -0.285 mm relative to bregma, DV measurements relative to the skull surface). Tips of the angled fiber optics were positioned on the lateral side of the target location. Cannulas were cemented in place with Metabond and Amalgambond Catalyst (Parkell) and animals were allowed to recover for at least 2 weeks prior to handling and water restriction.

#### Recording

Fiber photometry signals were recorded with a low autofluorescence fiber optic patch cord (Doric Lenses, MFP_400/430/1100-0.57_1m_FC-MF2.5_LAF) coupled to the animals with a brass mating sleeve. Photometry systems were captured using either the RZ5P or RZ10X LUX-I/O digital acquisition system. *RZ5P setup*. Two LED light sources (405 nm and 490 nm) were sinusoidally modulated by LED drivers at frequencies of 220 and 330 Hz, respectively. Light from LEDs was filtered through a custom optical path as described in Lerner et al. Cell 2015 (Figure 4). Fluorescence was captured by a photodetector (Newport) and processed with a RZ5P digital acquisition system and Synapse software (Tucker-Davis Technologies). *RZ10X LUX-I/O setup*. Two LED light sources (405 nm and 465 nm) were sinusoidally modulated by LED drivers at frequencies of 220 and 330 Hz, respectively. Light from LEDs was filtered through a fluorescence minicube (Doric Lenses, FMC6_IE(400-410)_E1(460-490)_F1(500-540)_E2(555-570)_F2(580-680)_S). Fluorescence was captured by integrated photosensors and processed with a RZ10X LUX-I/O digital acquisition system and Synapse software (Tucker-Davis Technologies). Photometry data from both systems was synced to behavioral data and task events using TTL outputs captured by the same system including trial starts, nose pokes, entries into the reward port, predictive cues, air puffs, and shocks. Photometry data was analyzed offline using MATLAB as described below (see *Quantification and Statistical Analysis*).

#### Histology

Following all experiments for each cohort, animals were deeply anesthetized using an intraperitoneal injection of Euthasol (MWI Animal Health) and transcardially perfused with phosphate-buffered saline (PBS) and 4% paraformaldehyde (PFA) and postfixed overnight at 4 °C. For fiber photometry experiments, brains were sectioned at 75 µm thickness near the site of cannula placements. Brains were permeabilized in 0.1% TX100 and incubated overnight in anti-GFP antibody (Thermo Fisher Cat # A-31852 or Cat # A-21311 at 1:2000 dilution) to stain GCaMP-expressing neurons. Sections were washed in 1xPBS and mounted on coverslips with Fluoromount-G. Brain sections were imaged on a Nikon ‘Sora’ Spinning Disk Confocal Microscope with 405 nm and 640 nm or 488 nm lasers and a 20x objective.

### Tac1 expression labeling

#### Surgical procedure

For labeling LHb^Tac1^ expression, 200 – 300 nL of AAV1-Ef1a-DIO-YFP was injected in the right or left LHb of Tac1-Cre male and female mice age 13-18 weeks and transcardially perfused 8 weeks post-op using the surgical procedure and perfusion protocol described above. Brains were sectioned at 75 µm thickness from ∼ 0 mm Bregma – ∼-4.8 mm Bregma. Brains were permeabilized in 0.1 % TX100 and incubated overnight in anti-GFP antibody (Thermo Fisher Cat # A-21311 at 1:2000 dilution) to stain YFP-expressing neurons. Sections were washed in 1xPBS and mounted on coverslips with Fluoromount-G. Brain sections were imaged on a Nikon ‘Sora’ Spinning Disk Confocal Microscope with 405 nm and 488 nm lasers and a 20x objective. Images were analyzed with a custom MATLAB app. Individual sections were assigned to an AP atlas position and cells inside the LHb boundaries were manually annotated. Next, 6-10 reference points were identified to register each section and its associate cells to the atlas image using an affine transformation and forward projection.

### Quantification and Statistical Analysis

#### Fiber Photometry

Fiber photometry data was collected using Synapse software (TDT instruments). Timestamps for behavioral epochs were collected from operant training boxes and synced with photometry data in Synapse Software. Briefly, for each behavioral session the 405 nm signal was fit to the GCaMP signal and subtracted from it. The resulting signal was downsampled to 20 Hz and reported as Normalized F. Z-scores were calculated using the Normalized F across the entire behavioral session. Comparisons across animals and genotypes were reported as z-score to reduce effects of variability in viral expression across animals. For experiments where event statistics would lead to a change in z-score across the data being compared; for example, in comparing across sessions of varying reward probabilities, we reported Normalized F.

#### Reward History Analysis

We used a linear mixed-effect model to predict trial-by-trial changes in fluorescence from a set of binary regressors indicating whether each of the 10 prior trials was rewarded or omitted. For the effect of prior size, we used reward size as a regressor over the previous 5 trials. Models were fit separately for each subject, with additional covariates for trial number, session reward probability, and prior choice history (correct, premature, or no response). To evaluate significance, we performed a permutation test in which the omission predictors were shuffled 1000 times to generate a null distribution of the coefficients and compared the median real beta coefficients to the null distribution of medians. P values were corrected for multiple comparisons using the Benjamini-Hochberg false discovery rate.

#### Trial Type Classification

We trained a multiclass error-correcting output code classifier (support vector machine) to predict either cue identity, delivered reward size, expected reward size, or RPE on each trial based on the trial-by-trial changes in the time series of fluorescence from 0-5 seconds after reward port entry with subject ID included as a predictor to account for variation in viral expression levels. For classifying cue identity, trial reward outcome (delivered or omitted) was also included as a predictor (Figure 2L). For joint classification of reward size and expectation, each expectation-size combination represented a unique class (Figure S3D). To compare models using LHb^Tac1^ activity to classify trial types by either size, expectation, or RPE (Figure 4G-J) we analyzed 10 µL and 30 µL reward delivery across 10 µL and 30 µL expected reward size. All models were trained using only correct trials from each session (premature and no response excluded) and were subsampled to the number of trials in the minority class to create balanced datasets within each model, with ten-fold cross-validation.

